# Exposure to Antibiotics Modifies the Immune Profiles of Bacterial Extracellular Vesicles from Common Vaginal Anaerobes

**DOI:** 10.64898/2026.05.21.726874

**Authors:** Yu Hasegawa, Olivia Swain, Urvija Rajpal, Michael France, Liqhwa Ncube, Ilaria Mogno, Hannah Zierden, Jacques Ravel, Michal A. Elovitz

## Abstract

**Background:** The female lower reproductive tract harbors a complex microbiome that plays a critical role in reproductive health. A vaginal microbiome dominated by *Lactobacillus crispatus* (LC; Community State Type (CST) I) supports vaginal health, whereas a microbiome enriched with anaerobic species, such as *Gardnerella vaginalis* (GV) and *Mobiluncus mulieris* (MM) (CST IV) is linked to bacterial vaginosis (BV) and adverse outcomes, including sexually transmitted infections, infertility, and preterm birth. Although antibiotics such as metronidazole and clindamycin are commonly prescribed to treat BV, recurrence rates remain high, and the impact of these treatments on bacterial extracellular vesicles (bEVs), critical mediators of host-microbe interactions, is poorly understood.

**Result:** We investigated how antibiotic treatment at a dose below minimum inhibitory concentration alters the production and immunomodulatory function of bEVs derived from GV, MM, and LC. Using nanoparticle tracking analysis, cytokine profiling, and TLR pathway analyses, we found that antibiotic treatment significantly enhanced the inflammatory properties of bEVs in a species- and antibiotic-specific manner. Notably, bEVs from antibiotic-exposed GV and MM cultures induced elevated cytokine responses in epithelial and immune cells, primarily through TLR2 activation for GV bEVs, and through both TLR2 and TLR5 activation for MM bEVs. While LC bEVs are typically non-inflammatory, exposure to metronidazole, even at a lower dose than what is used clinically, rendered them immunostimulatory, suggesting a potential unintended proinflammatory consequence of treatment on beneficial microbes. We also detected bEVs in human vaginal swabs, including vaginolysin-positive bEVs, even in CST I microbiomes, indicating that low-abundance microbes, including pathogens, remain transcriptionally active.

**Conclusions:** These findings suggest that antibiotics not only reduce microbial load but also reshape bacterial communication via bEVs, potentially contributing to inflammation, epithelial barrier disruption, persistent dysbiosis, and recurrent infections. This work underscores the need for precision antimicrobial strategies that eliminate pathogens while preserving beneficial bacteria and their functional bEVs. Future therapies may benefit from considering the ecosystem-wide effects of antibiotics on the vaginal microbiome and its bEV-mediated signaling network.

## Introduction

The female lower reproductive tract is a complex ecosystem comprising of host epithelial and immune cells, as well as microorganisms and the metabolites they produce ^1^. An optimal human vaginal microbiome is dominated by *Lactobacillus crispatus* (LC), categorized as Community State Type (CST) I ^2^. In contrast, microbiomes lacking *Lactobacillus* spp. and enriched with anaerobes such as *Gardnerella, Mobiluncus*, and *Prevotella* are classified as CST IV ^2^, which is associated with bacterial vaginosis (BV), as well as adverse reproductive outcomes such as infertility, endometriosis, preterm birth (PTB), and premature preterm rupture of membranes (PPROM) ^3,4^. Antibiotics are widely used in clinical care for these conditions to eliminate pathogenic anaerobes and reduce associated inflammation. BV, a common condition affecting 20-50% of reproductive age women, is characterized by symptoms such as greyish vaginal discharge, odor, itching, and dysuria ^5,6^. While BV is defined clinically, it is microbiologically characterized by a paucity of *Lactobacillus* spp. and the presence of a diverse array of strict and facultative anaerobes ^5^. Oral or intravaginal administration of metronidazole or clindamycin is commonly prescribed to alleviate BV symptoms and reduce the abundance of anaerobes such as *G. vaginalis*. However, the recurrence rates remain high, with up to 60-80% of patients experiencing relapse within 6-12 months after treatment ^5^. Although the development of antibiotic resistance is well-recognized, it is under-appreciated in BV treatment and the broader ecological impact of antibiotic therapy on the vaginal ecosystem remains poorly understood.

We have recently demonstrated that vaginal bacteria produce functional and unique bacterial extracellular vesicles (bEVs) that play a role in host-microbe interaction *in vitro* ^7^. BEVs are released by both gram-negative and gram-positive bacteria and encapsulate a wide range of bacterial bioactive molecules (*e*.*g*., nucleic acids, proteins, lipids, metabolites) that can be delivered to distant sites ^8,9^. BEVs mediate both bacteria-bacteria and bacteria-host interactions ^8^. The interaction between bEVs and antibiotics is an emerging area of study and likely involves complex mechanisms. These vesicles have been implicated in contributing to antibiotic resistance by acting as decoys that sequester antibiotics or by carrying enzymes that degrade them ^9^. Antibiotic exposure has also been shown to alter bEV biogenesis ^10^. However, despite the routine use of antibiotics to treat BV, there is a paucity of data on how antibiotics affect bEV production and function within the vaginal microbiome. Here, we sought to model a scenario in which antibiotic treatment fails to completely eradicate bacteria, allowing a subset of bacteria to survive while undergoing antibiotic□induced metabolic alterations. We hypothesized that sublethal dose of antibiotic exposure modulates bEV functionalities in ways that increase inflammation, potentially contributing to high recurrence rates and/or persistent symptoms in patients treated for BV. To investigate the effects of antibiotic exposure on bEV production and function, we assessed bEVs produced by two common pathogenic vaginal anaerobes, *G. vaginalis* (GV) and *M. mulieris* (MM), as well as a beneficial bacterium, LC. GV has been extensively studied as a key pathogen associated with adverse health and reproductive outcomes ^3,11^. Although MM is less abundant than GV, it shows a strong association with the risk of spontaneous PTB ^11–13^ and remains relatively understudied. We tested metronidazole and clindamycin, which are routinely prescribed to treat BV ^5,6,14^. To determine whether the impact on bEVs was specific to these antibiotics or if it reflected a broader, non-specific response, we also investigated ampicillin, a broad-spectrum antibiotic not typically used to target vaginal microbes but routinely prescribed to women for conditions such as urinary tract infections ^15,16^, pneumonia ^15^, as well as PPROM prophylaxis ^16,17^ and chorioamnionitis ^16^.

The study addressed three main questions: 1) whether antibiotic exposure alters the quantity of bEVs produced; 2) whether bEVs from antibiotic-exposed bacterial cultures elicit different immune responses in epithelial and immune cells compared to bEVs from untreated cultures; and 3) whether Toll-like receptor (TLR) pathways play in mediating the immune responses triggered by bEVs from vaginal anaerobes.

## Methods

### Cell culture conditions

Human ectocervical (Ect; American Type Culture Collection (ATCC) CRL-2614, Ect1/E6E7, Manassas, VA, USA), endocervical (End; ATCC CRL-2615, End1/E6E7), and vaginal (VK2; ATCC CRL-2616, VK2/E6E7) epithelial cell lines ^18^ were cultured in keratinocyte-serum-free medium supplemented with 0.1 ng/mL epidermal growth factor, 50 μg/mL bovine pituitary extract (Gibco/Thermo Fisher Scientific, Waltham, MA, USA), and 100 U/mL penicillin and 100 μg/mL streptomycin (Gibco). THP-1-Null cells (THP1) and THP1-Dual KO-TLR2 cells (THP1-TLR2KO; InvivoGen, San Diego, CA, USA) were cultured in RPMI-1640 medium (Life Technologies, Grand Island, NY, USA) supplemented with 10% (v/v) heat-inactivated fetal bovine serum (HI-FBS) (Gibco) and 100 U/mL penicillin and 100 μg/mL streptomycin. HEK-Blue reporter cell lines (HEK-Blue hTLR2, HEK-Blue hTLR2-TLR1, HEK-Blue hTLR2-TLR6; InvivoGen) were maintained in DMEM supplemented with 10% (v/v) HI-FBS, 100 U/mL penicillin and 100 μg/mL streptomycin, 100 μg/mL normocin (InvivoGen), and 1X HEK-Blue Selection (InvivoGen).

### Bacterial culture and sample preparation

All bacterial strains were grown in NYCIII medium supplemented with 1% horse serum (Gibco). The culture medium was pre-processed by ultracentrifugation at 100,000 × g overnight using an Optima Max-XP Tabletop Ultracentrifuge (Beckman Coulter, Jersey City, NJ, USA) to remove any potentially contaminating EVs. Clinical strains of LC (ATCC 33197), GV (ATCC 14018), and MM (ATCC 35243) were cultured in 300 mL of NYCIII medium, either without antibiotics or with varying doses of ampicillin (A9518-5g, Sigma-Aldrich, St. Louis, MO, USA), clindamycin (HY-B0408A/CS2508, MedChemExpress, Monmouth Junction, NJ, USA), or metronidazole (M1547-5G, Sigma-Aldrich) to determine the minimum inhibitory concentration (MIC) and the required growth duration. To isolate bEVs for further *in vitro* experiments, the three bacterial strains were cultured at the predetermined sublethal MIC (Sub-MIC) at 37°C in an anaerobic glove box for a predetermined length of incubation (**Table 1**).

**Table 1.** Antibiotic doses and the duration of bacterial culture used in this study. The antibiotic doses in the table represent Sub-MIC for each condition, and the days in parentheses indicate the length of bacterial culture used to collect bacterial supernatant for bEV isolation.

|  | Ampicillin | Clindamycin | Metronidazole |
| --- | --- | --- | --- |
| *Lactobacillus crispatus*<br>(LC) | 1 ug/mL<br>(7 days) | 0.01 ug/mL<br>(7 days) | 512 ug/mL<br>(7 days) |
| *Gardnerella vaginalis*<br>(GV) | 0.05 ug/mL<br>(2 days) | 0.01 ug/mL<br>(3 days) | 8 ug/mL<br>(4 days) |
| *Mobiluncus mulieris*<br>(MM) | 0.05 ug/mL<br>(10 days) | 0.01 ug/mL<br>(10 days) | 2 ug/mL<br>(10 days) |

The bacterial culture medium was first centrifuged at 3,500 × *g* for 30 minutes, followed by filtration through a 0.02 µm Nalgene vacuum filtration system (Fisher Scientific, Fair Lawn, NJ, USA) to remove live bacteria and cell debris. BEVs were isolated from the filtrate by sequential ultracentrifugation and quantified as previously described ^7^. Briefly, the supernatant was centrifuged at 30,000 × *g* for 33 minutes to remove any remaining particles. The supernatant was then ultracentrifuged at 100,000 × *g* for 70 minutes to collect the pellets, which were washed once with phosphate-buffered saline (PBS) and centrifuged again at 100,000 × *g* for 70 minutes. The final bEV pellets were resuspended in EV suspension medium (composed of 10 mM HEPES (Gibco) and 25 mM NaCl solution) and stored at -80°C until further analysis. The quantity and size distribution of BEVs were determined using Nanoparticle Tracking Analysis (NTA) with ZetaView (Particle Metrix, Germany).

### Treatment of cells with bEVs

To optimize the bEV dose per well, bEVs isolated from GV and MM cultures that were not exposed to any antibiotics were added to 1.5 × 10^5^ End1 cells at 10^7^, 10^8^, and 10^9^ and measured GM-CSF, IL-6, and IL-8 levels produced by cells in the cell culture supernatant by Luminex (further details are stated below). As a result, while 10^7^ bEVs were sufficient to induce significant elevations in all mediators tested for GV bEVs, 10^8^ bEVs were required to induce significant elevation for MM bEVs (Supplementary Figure 1). Therefore, for all bacteria and experiments, we used 10^8^ bEVs per well in this study.

Ect1, End1, VK2, and THP1 cells from three different passages were plated at 1.5 × 10^5^ cells per well in 24-well plates. After 24 hours, cells were cultured in growth medium (Media control), growth medium containing the same concentration of bEV suspension medium (10 mM HEPES (Gibco) and 25 mM NaCl) as used in other bEV treatment groups (bEV vehicle control), or with bEVs isolated from the supernatant of bacteria cultured without any exposure to antibiotics (Unexposed) or with antibiotic treatment (ampicillin, clindamycin, metronidazole) at 10^8^ bEVs per well. For dose-response tests, cells were treated with 10^7^ or 10^9^ bEVs per well. Because the bEV yield for the metronidazole group of LC cultures was too low to conduct the experiment under the same conditions as the other groups, a 96-well plate was used, ensuring that the cell-to-bEV ratio and the culture duration remained consistent with the other treatments. After 24 hours of incubation with treatment, the cell culture supernatant was collected and stored at -80°C until further analysis. To investigate the involvement of TLR5, THP1-TLR2KO cells were pre-treated with anti-hTLR5-IgA2 (InvivoGen) at a final concentration of 1 µg/mL for 1 hour, followed by bEV MM treatment at the same dose and conditions as described above for 24 hours.

### Profiling of inflammation markers

Frozen cell culture supernatant samples were thawed at 4°C overnight, vortexed briefly, and centrifuged at 10,000 × *g* for 10 minutes. The collected supernatant was analyzed using Luminex technology with the Human Cytokine/Chemokine/Growth Factor Panel A kit (HCYTA-60K-06, ThermoFisher Scientific) to measure the following inflammation markers: Interleukin (IL)-8, IL-6, IL-1β, IL-18, granulocyte-macrophage colony-stimulating factor (GM-CSF), and tumor necrosis factor-alpha (TNF-α). The same three samples were loaded onto each plate to account for plate batch effects, and cell culture media were assessed to ensure that no background noise interfered with the data. IL-8 levels in the cell culture media from THP1 and THP1-TLR2KO cells were assessed using an ELISA kit (R&D Systems, Minneapolis, MN, USA)

### Cell reporter assay for TLR2 activity assessment

TLR2, TLR2/1, and TLR2/6 activation were measured using HEK-TLR2 reporter cell lines according to the manufacturer’s protocol. Briefly, 20 µL of bEV samples (3.3 × 10^7^ bEVs), positive controls (IL-1β at 1 ng/mL for TLR2, Pam3CSK4 at 10 ng/mL for TLR2/1, or fibroblast-stimulating lipopeptide-1 (FSL-1) at 1 ng/mL for TLR2/6), or negative control (HEK cell growth medium without antibiotics) was added to each well of a 96-well plate. Following this, 50,000 cells suspended in 180 µL of HEK-Blue Detection medium were added to each well. The plate was incubated at 37°C in 5% CO□ for 6 hours in the dark, and the level of secreted embryonic alkaline phosphatase was measured at an optical density (OD) of 655 nm using a spectrophotometer.

### Nanoimager (ONI) analysis on the EVs from human swabs

EVs were isolated from human vaginal swabs collected as part of the M&M study, where the bacterial taxa composition of these swabs was known and previously reported ^12^. Informed consent was obtained in writing from all participants, and approval was granted by the Institutional Review Boards (IRBs) at the University of Pennsylvania (IRB #818914) and the University of Maryland School of Medicine (HP-00045398).

To extract protein, each Dacron swab was placed in PBS containing a Pierce Protease Inhibitor tablet (ThermoFisher Scientific) and incubated for 5 minutes. The swab was then thoroughly agitated in the solution. The isolation media was centrifuged at 10,000 × *g* for 5 minutes, and the resulting supernatant was collected. EVs were isolated and stored as described earlier.

The EVs were immunolabeled using the EV Profiler 2 kit (ONI, San Diego, CA, USA) with anti-mouse Vaginolysin antibody (Absolute Antibody, Shirley, MA, USA), anti-human tetraspanin trio (CD81/CD63/CD9), and PanEV (ONI) as the EV control. The samples were imaged using the ONI instrument according to the manufacturer’s instructions. Single-EV levels were then quantified using the CODI platform (https://alto.codi.bio). A density-based clustering analysis with drift correction and filtering was performed to evaluate each vesicle.

### Statistical analysis

A Pearson correlation test was used to assess the correlation between total bEV particles and bacterial OD at the time of supernatant collection for bEV isolation. For the multiplex immunoassay results, data from different plates were adjusted using plate batch controls, and concentrations below the minimum detection concentration (MinDC) were substituted with MinDC/√2. A one-way ANOVA followed by Sidak’s multiple comparison test with false discovery rate (FDR) correction was used to compare group differences. To compare IL-8 levels between THP1 and THP1-TLR2KO cells, a two-way ANOVA with Sidak’s multiple comparison test was applied. HEK assay results were normalized to the average reading of the Media control group, followed by a one-way ANOVA with Tukey’s multiple comparison test. For ONI data analysis, the ratio of vaginolysin-EV in bEVs isolated from pure GV culture was used to normalize the ratio of vaginolysin-EV in the swab samples. Statistical significance was set at p-values < 0.05.

## Results

### Nanoparticle Tracking Analysis on bEVs from bacteria with and without antibiotic treatment

Optical Density (OD) of LC, GV, and MM cultures with and without various doses of ampicillin, clindamycin, and metronidazole were measured every 24 hours over 96 hours to determine the Sub-MIC (Supplementary Figure 2). Based on these results, the Sub-MIC values used for treatment were as follows: 1, 0.01, and 512 µg/mL for LC; 0.05, 0.01, and 8 µg/mL for GV; and 0.05, 0.01, and 2 µg/mL for MM for ampicillin, clindamycin, and metronidazole treatments, respectively (Table 1). According to NTA, antibiotic treatments tended to reduce bEV concentrations compared to the Unexposed group (Table 2, **Figure 1**). The number of total bEV particles and the OD of bacterial culture media were not significantly correlated (p = 0.77, r = 0.095).

**Table 2.** BEV concentrations, as well as estimated bacterial growth curve and OD at the time of bacterial supernatant collection.

| Bacteria | ABX treatment | BEV concentration<br>(particle/mL) | Average bacterial<br>culture OD |
| --- | --- | --- | --- |
| LC | Unexposed | 9.30E+10 | 0.10 |
|  | Ampicillin | 3.50E+10 | 0.09 |
|  | Clindamycin | 1.00E+11 | 0.09 |
|  | Metronidazole | 5.20E+09 | 0.11 |
| GV | Unexposed | 1.90E+11 | 0.17 |
|  | Ampicillin | 2.00E+10 | 0.22 |
|  | Clindamycin | 2.50E+11 | 0.09 |
|  | Metronidazole | 1.90E+11 | 0.24 |
| MM | Unexposed | 6.80E+10 | 1.20 |
|  | Ampicillin | 3.50E+10 | 0.57 |
|  | Clindamycin | 4.10E+10 | 1.28 |
|  | Metronidazole | 4.40E+10 | 0.23 |

**Figure 1.**
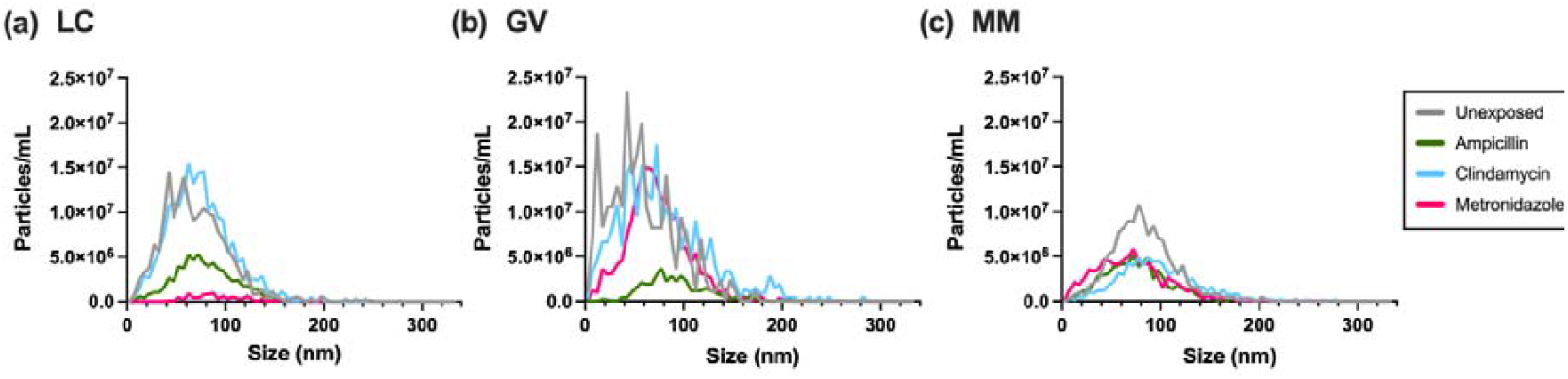
Size distribution of bEVs with and without antibiotic exposure. The bEV size distribution was visualized for bEVs isolated from (a) LC, (b) GV, and (c) MM. Grey lines represent bEVs isolated from bacterial cultures without any exposure to antibiotics (Unexposed) or treated with Sub-MIC of ampicillin (green), clindamycin (blue), or metronidazole (red) (n=1).

### Antibiotic treatments significantly elevated immunogenic properties of bEVs in bacteria- and antibiotic-specific manner

The Luminex multiplex immunoassay was used to measure the inflammatory mediators (GM-CSF, IL-6, IL-8, IL-1β, IL-18, and TNF-α) produced by cervical and vaginal epithelial cells, as well as THP1 monocytes.

LC bEV from both unexposed and antibiotic-exposed cultures did not significantly affect any inflammatory mediators in Ect1 and End1 cells. However, bEVs from LC exposed to metronidazole tended to increase IL-8 level, which did not reach statistical significance after FDR correction due to the large variability (**Figures 2a-2c**). In VK2 cells, while bEVs from unexposed cultures did not significantly alter any mediators’ levels, bEVs from LC exposed to metronidazole significantly increased all immune mediators compared to the bEV vehicle control group (IL-6, IL-8) and/or the Unexposed group (GM-CSF, IL-6) (Figures 2a-2c). Other antibiotic treatments did not result in a significant alteration in any mediator levels in comparison to the Unexposed group for LC.

**Figure 2.**
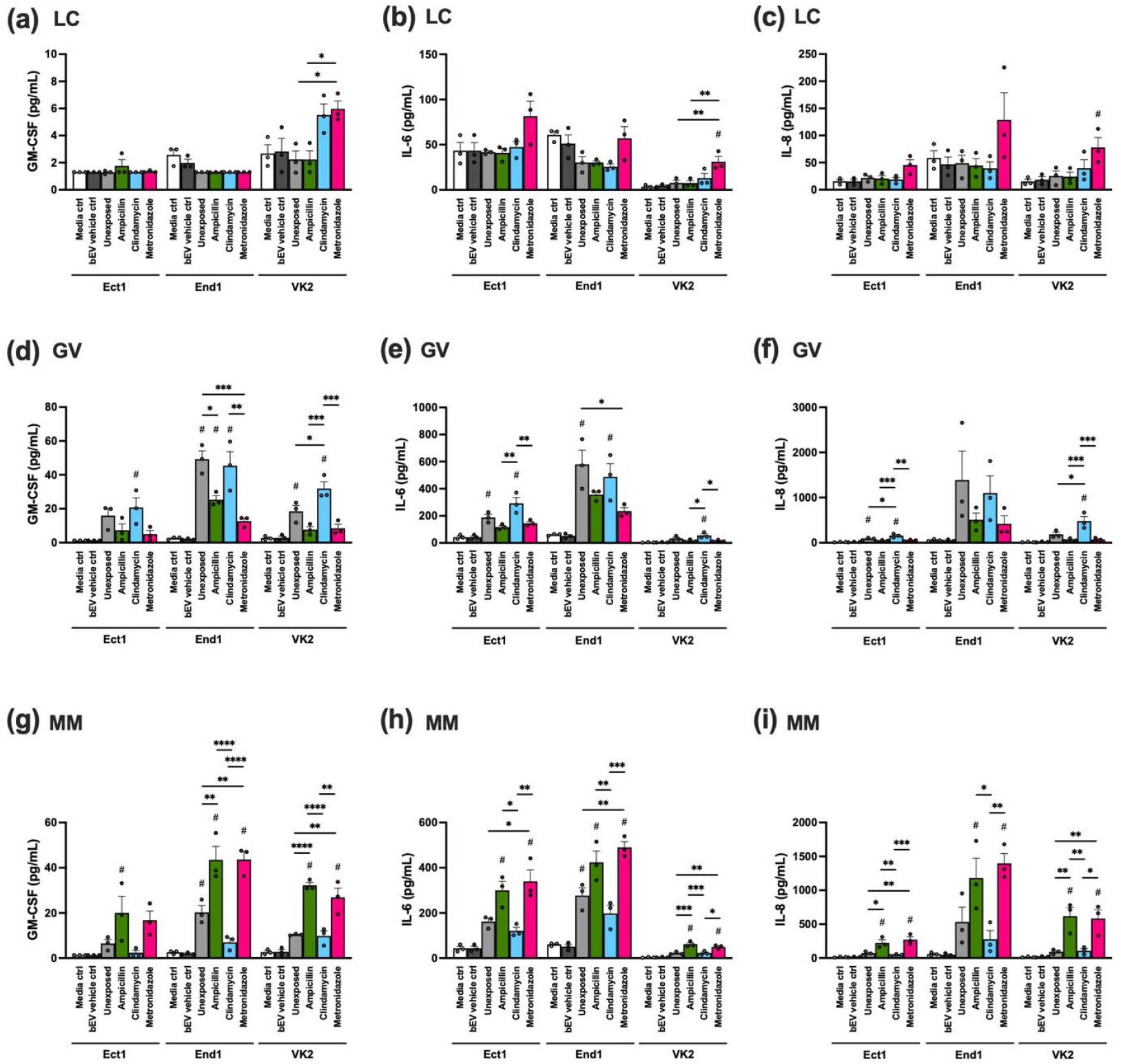
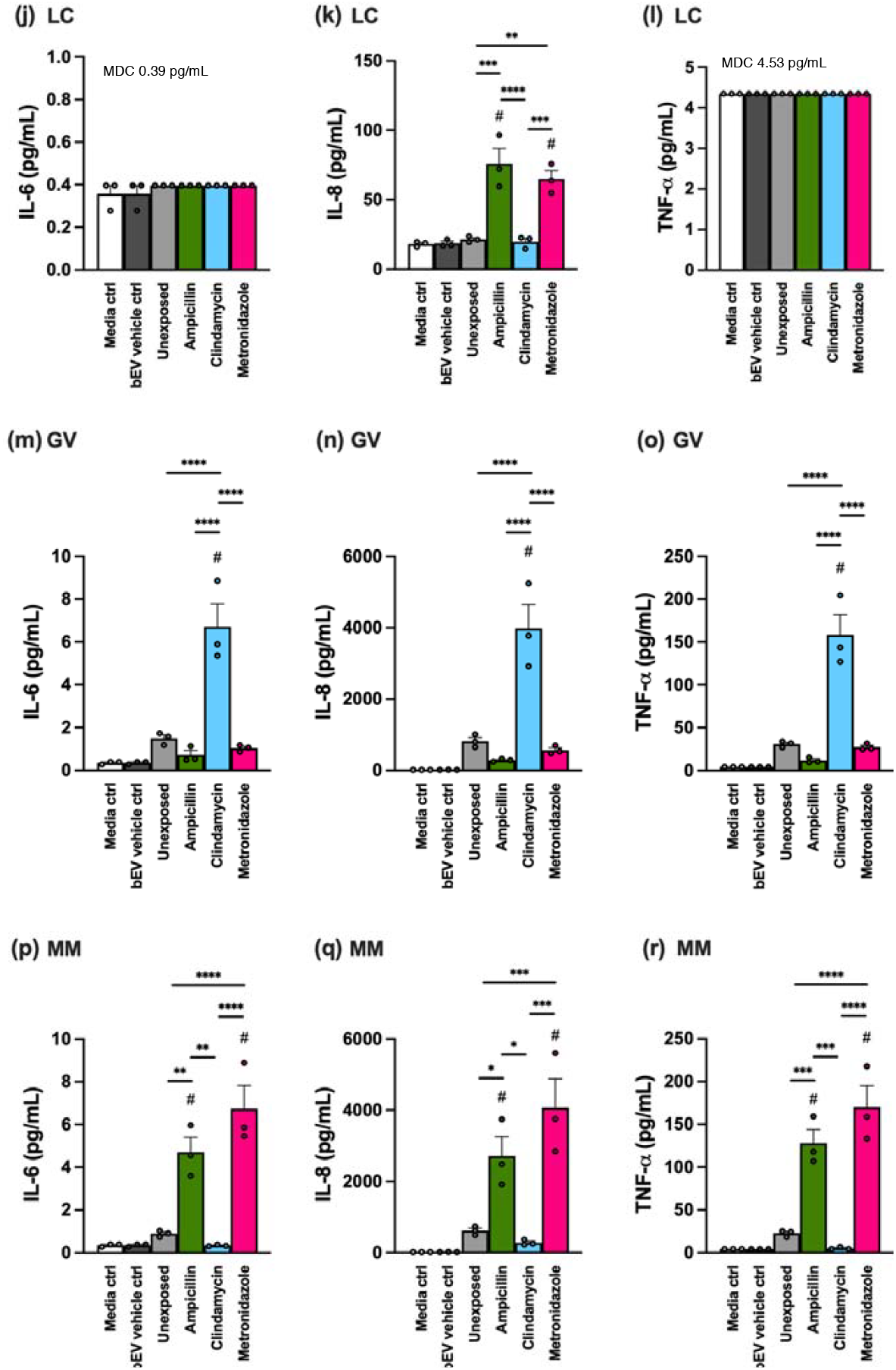
Production of cytokines by epithelial cells and monocytes in response to bEVs. We exposed ectocervical (Ect1), endocervical (End1) and vaginal (VK2) epithelial cells, as well as monocytes (THP1) with bEVs produced by LC, GV, and MM under different antibiotic treatment regimens: unexposed (grey), ampicillin (green), clindamycin (blue), metronidazole (red). As negative controls, cells were cultured either in growth media alone (Media ctrl, white) or in growth media containing the same concentration of bEV suspension media (bEV vehicle ctrl, dark grey). The levels of GM-CSF, IL-6, and IL-8 in the culture supernatant were measured in response to (a-c) LC bEVs, (d-f) GV bEVs, and (g-i) MM bEVs in Ect, End, and VK2 cells, respectively. Similarly, IL-6, IL-8, and TNF-α levels were measured in THP1 cells in response to (j–l) LC bEVs, (m–o) GV bEVs, and (p–r) MM bEVs. Minimal Detection Concentration (MDC) for IL-6 and TNF-α are included in the plot. Bar plots represent the mean levels, with error bars indicating the standard deviation (n=3). A one-way ANOVA followed by Sidak’s multiple comparison test with FDR correction was used to compare group differences. Significant differences from the bEV vehicle control group are indicated with “#,” and significant differences between bEV-treated groups are indicated with “*” (p<0.05), “**” (p<0.01), “***” (p<0.001), or “****” (p<0.0001).

As expected from our prior work ^7^, GV bEVs from unexposed cultures resulted in an increase in several inflammatory mediators (**Figures 2d-2f**). Importantly, exposure to antibiotics altered GV bEVs’ induction of the immune response. Compared to the Unexposed group, clindamycin exposure resulted in a significant elevation in IL-8 level in Ect1, as well as GM-CSF and IL-8 in VK2 cells. In contrast, ampicillin and/or metronidazole exposure resulted in a significant reduction in GM-CSF level in End1 cells in comparison to GV bEV unexposed to antibiotics. GV bEVs from cultures exposed to different antibiotics had varying effects on the ability of GV bEVs to induce an inflammatory response.

MM bEVs from unexposed cultures induced an immune response, with significant elevations in GM-CSF and IL-6 levels in End1 cells, whereas no significant alterations were found in Ect1 and VK2 cells (**Figure 2g-2i**). Compared to the Unexposed group, MM bEVs that were exposed to ampicillin significantly elevated IL-8 levels in Ect1, GM-CSF in End1, and all three mediators in VK2 cells. Additionally, metronidazole exposure resulted in significant elevation in IL-6 and IL-8 levels in Ect1, GM-CSF, and IL-6 levels in End1, and all three mediators in VK2 cells in comparison to the Unexposed group. Clindamycin exposure did not modulate the immunomodulatory properties of MM bEVs compared to the levels found in the Unexposed groups. Antibiotic exposure altered the ability of MM bEVs to induce inflammation from epithelial cells, but notably not in the same manner as observed for GV bEVs.

In THP-1 cells, compared with the bEV vehicle control group, the Unexposed group treated with bEVs from GV and MM showed increased levels of IL-6 (4.2-fold increase for GV, 2.5-fold increase for MM), IL-8 (44.0-fold increase for GV, 33.1-fold increase for MM), and TNF-α (7.1-fold increase for GV, 5.2-fold increase for MM).

However, these differences did not reach statistical significance after FDR correction for multiple comparisons (adjusted p-value > 0.05) (**Figures 2j-2r**). IL-6 and TNF-α levels by THP1 cells in response to LC bEVs were at minimal detection limit (MDC) (**Figure 2j**). In comparison to the Unexposed groups, bEVs from LC with ampicillin and metronidazole exposure significantly elevated IL-8 production (**Figure 2k**). In contrast, bEVs from GV and MM elevated all mediators to a higher degree compared to bEVs from LC. BEVs from GV with clindamycin exposure resulted in significant elevations in IL-6, IL-8, and TNF-α levels (**Figures 2m-2o**),and bEVs from MM with ampicillin and metronidazole exposure significantly elevated IL-6, IL-8, and TNF-α levels (**Figures 2p-2r**). The levels of IL-1β, IL-18, and TNF-α did not change by any exposure groups in Ect1, End1, and VK2 cells. Also, GM-CSF, IL-1β, and IL-18 were below the detection limit in THP1 cells.

### TLR2 and TLR5 are involved in mediating augmented immune responses by bEVs from antibiotic-exposed vaginal bacterial cultures

To assess whether TLR2 activation mediates the augmented immune response from bEVs produced in the antibiotic-exposed GV and MM cultures, THP1-TLR2KO cells were used, and IL-8 level was measured as an indicator of inflammation. While increased IL-8 from metronidazole-exposed LC was unexpected, we found that absence of TLR2 limited this induction of IL-8 (**Figure 3a**). Absence of TLR2 significantly reduced the ability of all GV bEVs (both Unexposed and antibiotics-exposed groups) to induce IL-8, bringing levels back to the level found in the bEV vehicle control group (**Figure 3b**). In contrast, IL-8 production in the Unexposed MM bEV group was significantly reduced, decreasing from 601.9 ± 14.2 pg/mL to 24.4 ± 4.0 pg/mL, and reaching levels close to those of the bEV vehicle control group (**Figure 3c**). However, the reduction caused by the absence of TLR2 in antibiotic-exposed MM bEV groups was significant, but the levels did not reach those found in the bEV vehicle control group (1,725.0 ± 60.1 to 302.3 ± 29.2 pg/mL in ampicillin group; 228.5 ± 5.7 to 111.1 ± 11.5 pg/mL in clindamycin group; 2,214.2 ± 39.8 to 114.6 ± 14.7 pg/mL in metronidazole group). Considering that MM possesses flagella, which are recognized by TLR5 (the receptor for flagellin) ^19^, we investigated the effect of a TLR5 inhibitor (TLR5i) (**Figure 3d**). Under these conditions, the IL-8 production induced by all MM bEV groups was significantly suppressed and returned to the levels found in the media control group.

**Figure 3.**
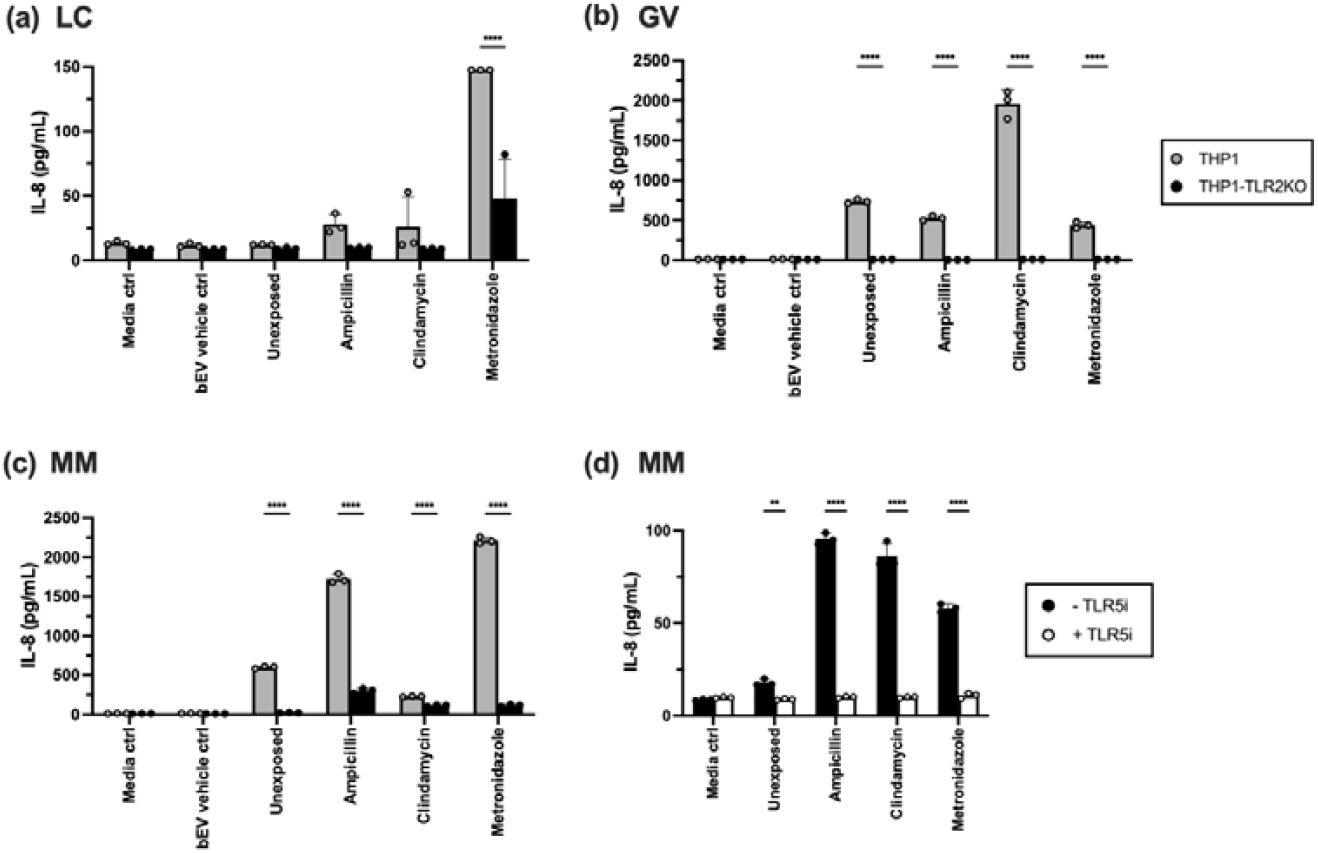
The involvement of the TLR pathway as the recognition pathway for bEV treatment. IL-8 levels in the supernatants of THP1 or THP1-TLR2KO cells were measured in response to bEVs isolated from (a) LC, (b) GV, and (c) MM, either without any exposed to antibiotics (Unexposed) or with antibiotics (ampicillin, clindamycin, or metronidazole). As negative controls, cells were cultured in growth media (Media ctrl) or in growth media with the same concentration of bEV suspension media (bEV vehicle ctrl). Bar plots represent the mean, with error bars indicating the standard deviation (n=3). Grey bars represent results from THP1 cells, and black bars represent results from THP1-TLR2KO cells. (d) IL-8 levels from THP1-TLR2KO cell supernatants treated with MM bEVs, with or without the TLR5 inhibitor (TLR5i). Black bars represent results from THP1-TLR2KO cells without TLR5i, and white bars represent results with TLR5i. Two-way ANOVA was used to assess the impact of bEV treatment, followed by Sidak’s multiple comparison test with FDR correction, with statistical significance denoted by “**” (p < 0.01) and “****” (p < 0.0001).

### BEVs from unexposed and antibiotic-exposed bacterial cultures activate specific TLR2 heterodimers

Since TLR2 consists of two types of dimers (TLR2/1 and TLR2/6) ^20^, we utilized HEK reporter cell lines to assess the specific impact of bEVs on each TLR2 dimer. For LC bEVs, while the Unexposed group did not induce an immune response in cervicovaginal epithelial cells and THP1 cells, it significantly increased total TLR2 activity (**Figure 4a**); however, the magnitude of this response was substantially lower than that observed with GV bEVs (**Figure 4b**) and MM bEVs (**Figure 4c**). On the other hand, the total TLR2 activity was significantly elevated by the Unexposed and all three antibiotics-exposed GV bEVs groups. While the activity of TLR2/1 was not significantly altered by any antibiotic-exposed groups, all groups significantly elevated that of TLR2/6. The increase in total TLR2 activity in the GV bEVs ampicillin group was significantly lower than that observed in the unexposed and other bEV-treated groups. Similarly to GV bEVs, all MM bEV groups significantly elevated total TLR2 activity with both TLR2/1 and TLR2/6 involved in the response, except for the clindamycin group, which was activated solely by TLR2/6. The elevation in total TLR2 activity by the clindamycin group was significantly lower than in the Unexposed and other bEV-treated groups.

**Figure 4.**
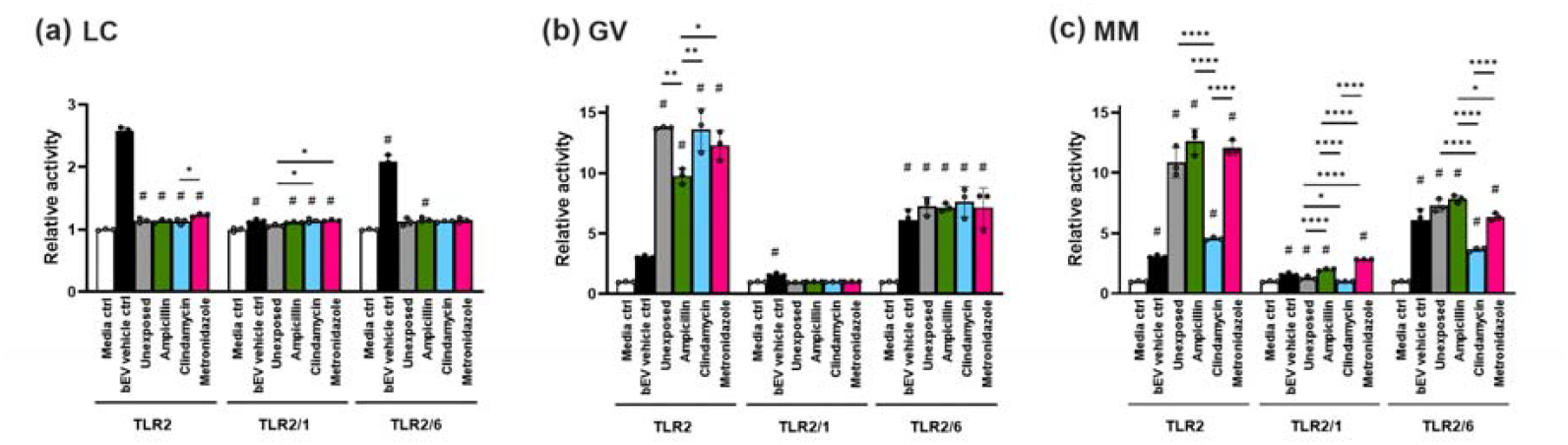
HEK assay measuring the relative activity levels of TLR2, TLR2/1, and TLR2/6 in HEK reporter cells in response to bEVs. HEK cells were exposed to bEVs produced by (a) LC, (b) GV, and (c) MM under different antibiotic treatment regimes: unexposed (grey), ampicillin (green), clindamycin (blue), metronidazole (red). As negative controls, cells were cultured either in growth media alone (Media ctrl, white) or in growth media containing the same concentration of bEV suspension media (bEV vehicle ctrl, dark grey). Raw readings were normalized to the average reading of the Media ctrl group. Bar plots represent the mean, with error bars indicating the standard deviation (n=3). A one-way ANOVA followed by Sidak’s multiple comparison test with FDR correction was used to compare group differences. Significant differences from the bEV vehicle control group are indicated with “#,” and significant differences between bEV-treated groups are indicated with “*” (p<0.05), “**” (p<0.01), “***” (p<0.001), or “****” (p<0.0001).

### BEVs from common vaginal anaerobes are present in the human vagina

To confirm that the human vagina contains bacterial EVs, EVs were isolated from six human vaginal swabs. Vaginolysin, a protein produced by GV ^21^ and encapsulated in GV-derived EVs ^7^, was targeted as a marker for bacterial EVs (vaginolysin-EV), while tetraspanin was used to identify host-derived EVs (Human-EV). The swabs were collected from two participants with CST I-A (ID526, ID532), three participants with CST IV-B (ID419, ID471, ID545), and one participant with CST III-B (ID1024) (**Figure 5a**). The vaginal microbiota composition of these samples varied: ID526 and ID532 were composed of 99.5% and 97.5% LC, respectively, with total GV of only 0.000008% and 0.028%. In contrast, ID419, ID471, ID545, and ID1024 had total GV levels of 22.4%, 27.0%, 23.1%, and 21.6%, respectively. Based on the ONI analysis, both vaginolysin-EV (**Figure 5b**) and human-EV (**Figure 5c**) were detected in all six human swab samples. The ID526 and ID532 samples showed 38% and 86% of vaginolysin-EV, respectively (**Figure 5d**). Notably, the percentages of vaginolysin-EV detected did not correlate with the total GV percentage (p = 0.50, r = -0.35).

**Figure 5.**
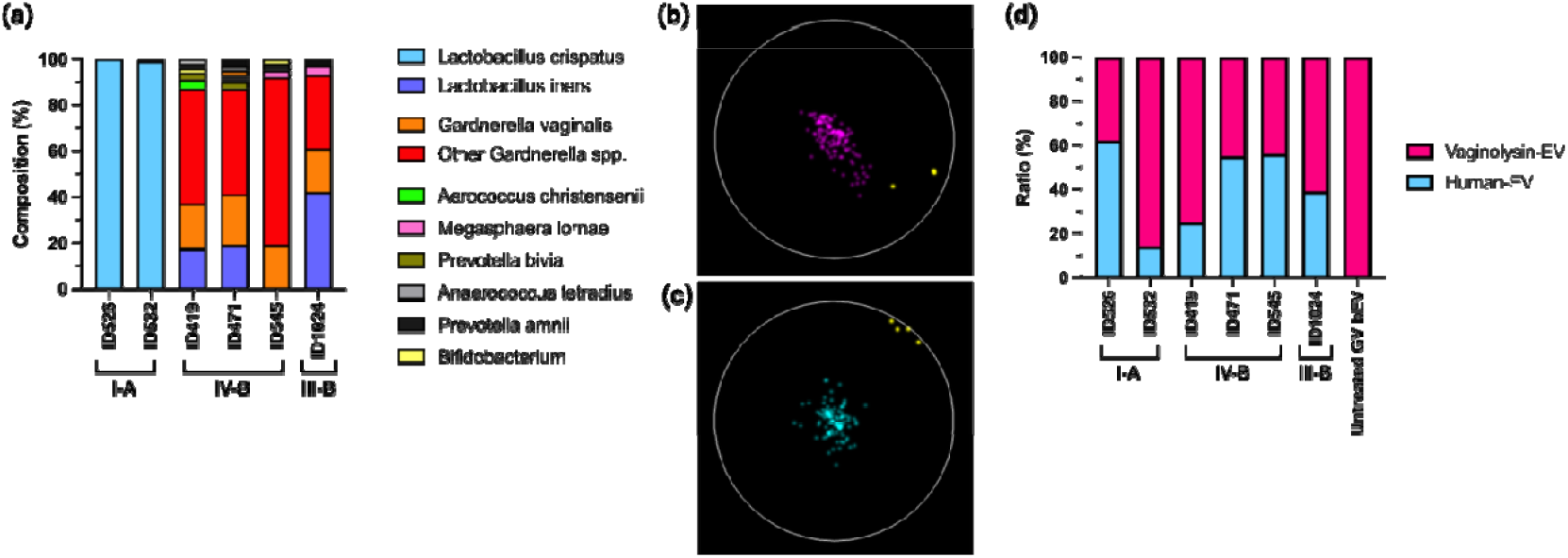
Presence of EVs from human and bacterial origins. (a) Bacterial taxa composition in vaginal swab samples. The color scheme is as follows: LC is light blue, *L. iners* is dark blue, GV is orange, other Gardnerella species are red, and other abundant microbial taxa are shown in the legend. The % composition of MM is too low to be visible in the plot. EVs were visualized using the Nanoimager system, targeting (b) vaginolysin (Vaginolysin-EV; red) and (c) tetraspanin trio (Human-EV; blue). EV markers are visualized in yellow. (d) Ratio of EVs positive for vaginolysin relative to EVs classified as human-derived in each vaginal sample. The ratio of Vaginolysin-EV in each sample was normalized to the ratio found in bEVs isolated from pure Untreated GV cultures.

## Discussion

The microbiome and microbial-host interactions in the cervicovaginal space are increasingly recognized as critical factors in reproductive health and disease. Yet, the complexity of this ecosystem is only beginning to be unraveled. We have recently demonstrated that several of the most common vaginal anaerobes produce bEVs similar to those observed in gram-positive microorganisms in other biological niches ^7^. Given the known impact of antibiotics on bEV biogenesis and function ^9,10^ and the widespread use of antibiotics to treat clinical conditions associated with the vaginal microbiome ^6^, we sought to determine the effect of commonly used antibiotics on the production and function of bEVs from bacteria associated with reproductive health and disease. The novel findings presented herein demonstrate a significant impact of antibiotics on the function of bEVs from these prevalent vaginal anaerobes. Notably, antibiotics commonly used to decrease colonization by pathogenic anaerobes in the cervicovaginal space led to the production of bEVs that increase inflammation in both epithelial and immune cells. Importantly, we reveal specific TLR-mediated mechanisms by which these bEVs drive local inflammation.

Antibiotic treatment at a dose below the MIC were able to alter the immunomodulatory properties of bEVs isolated from GV and MM culture in a manner that was specific to the bacterial species, host cell line, and type of antibiotics used. While exposure to antibiotics, particularly ampicillin and metronidazole, tended to reduce host inflammatory response induced by GV bEV, these antibiotics significantly altered MM bEV, resulting in a significant elevation of inflammatory responses by host cells. One potential mechanism by which antibiotic treatment may exacerbate host inflammation is by increasing bEV release. Previous studies have shown that antibiotic exposures can enhance bEV secretion compared to levels observed under non-exposed conditions ^22,23^, potentially as a bacterial survival strategy in response to antibiotic pressure ^10^. However, in this study, we did not observe a significant increase in bEV production following antibiotic exposure (Table 2). Notably, Torabian *et al*. reported that b-lactam antibiotics significantly increased bEV release from *Escherichia coli*, while quinolone and aminoglycoside antibiotics caused reduced vesiculation ^22^. These findings suggest that the impact of antibiotic exposure on bEV production likely depends on multiple factors, including the type of antibiotic, its mechanism of action, concentration, treatment duration, as well as the bacterial strain, culture conditions and growth phase ^22,23^. In our study, antibiotic exposures likely altered the immunomodulatory properties of bEVs by modifying their composition, thereby influencing their biological function and impact on the host cells.

Although all three bacterial strains were exposed to the same concentration, clindamycin significantly exacerbated the immunomodulatory property of only GV bEVs in Ect1, VK2, and THP1 cells compared to bEVs isolated from GV culture without any exposure to antibiotics. Clindamycin prevents bacterial growth by inhibiting bacterial protein synthesis. It binds to the 50S ribosomal subunit ^24^, preventing peptide chain elongation during translation by interfering with aminoacyl-tRNA transfer ^24^, thereby inducing metabolic stress. Consequently, the observed alteration in GV bEVs following clindamycin exposure may be primarily due to changes in GV protein content. In contrast, ampicillin and metronidazole significantly elevated the immunomodulatory properties of MM bEVs across all cell lines. Ampicillin is a b-lactam antibiotic in the penicillin class ^16^. It disrupts bacterial cell wall synthesis by binding penicillin-binding proteins (PBPs), which are essential for the final steps of peptidoglycan cross-linking ^16^. This disruption weakens the bacterial cell wall, resulting in abnormalities in cell shape, impaired division, cell lysis, and, ultimately, bacterial death ^16,25^. Moreover, PBP inhibition can interfere with the anchoring of flagellar machinery to the peptidoglycan layer, thereby impairing flagellar motility, localization, and function ^25,26^. Metronidazole, a nitroimidazole antibiotic ^14^, is activated in anaerobic environments, where its nitro group is reduced by bacterial enzymes to generate reactive nitrogen species ^27^. These species induce DNA strand breaks and cell stress, leading to bacterial cell death ^27^. Therefore, the elevated immunomodulatory activity of MM bEV in response to ampicillin and metronidazole may be attributed to changes in their cell wall and stress. To better understand how antibiotic exposure differentially affects bEV composition and the resulting immune responses in host cells, further proteomic analyses of bEVs derived from antibiotic-exposed bacteria are required.

Our findings revealed that ligands recognized by TLR signaling pathways are present in bEV cargo and that immune responses mediated by TLR2 dimer activation exhibit bacterial- and antibiotic-specific differences. TLR2 recognizes bacterial lipoproteins through two major heterodimer partners: TLR2/6, which binds diacylated lipoproteins typically found in gram-positive bacteria, and TLR2/1, which binds triacylated lipoproteins commonly found in gram-negative bacteria ^20^. Both dimers trigger the same downstream signaling cascade via MyD88, leading to NF-κB activation and subsequent induction of innate immune functions such as cytokine production, phagocytosis, and antimicrobial responses ^20^. We observed that GV bEVs were primarily recognized by TLR2, which is consistent with our previous findings ^7^, specifically through the TLR2/6 dimer. Interestingly, although clindamycin exposure resulted in greater induction of inflammatory mediators compared to other GV bEV groups, this was not mirrored by increased TLR2 activity, suggesting that additional pattern recognition receptors may be involved in detecting clindamycin-altered bEVs. In contrast, MM bEVs were also recognized by TLR2, as previously observed ^7^, with both TLR2/1 and TLR2/6 dimers being activated, except in clindamycin-exposed group, which was recognized solely by TLR2/6. This suggests a more complex lipoprotein profile in MM bEVs than GV bEVs. Notably, all three antibiotics exposure altered MM bEVs in a way that also activated TLR5, suggesting structural modifications to the bacterial components, particularly the cell wall and protein compositions, leading to changes in the component and abundance of flagellin in MM bEVs. The antibiotic-driven modulation of bEV recognizability by host TLRs adds to the complexity of the cervicovaginal microbial ecosystem and may present opportunities for therapeutic strategies to minimize the inflammatory impact of pathogenic bacteria and their bEVs.

Importantly, we also examined the impact of antibiotic exposure on health-promoting *L. crispatus*. As we previously reported ^7^, LC bEVs did not induce a significant immune response under unexposed conditions. However, we found that a high dose of metronidazole altered LC bEVs in a way that induced an immune response in both epithelial and immune cells. Notably, the concentration of metronidazole that elicited this pro-inflammatory response (512 µg/mL) is much greater than what would present in the vagina during clinical metronidazole treatment (vaginal metronidazole concentration ranged between 24 to 49 µg/mL after oral intake of 200 mg for 2 days, followed by 800 mg for 5 days, three times a day ^28^).

Therefore, metronidazole itself would not be likely to adversely affect LC clinically. However, our study showed that exposure to sublethal doses of all three commonly used antibiotics led to the release of bEVs from anaerobic bacteria commonly found in the vaginal environment associated with CST IV. These bEVs induced significantly increased pro□inflammatory cytokine production in cervicovaginal epithelial cells. Such inflammatory environment can become hostile to beneficial *Lactobacillus* spp., impairing their colonization and potentially favoring the growth of anaerobes. Such a shift may undermine the therapeutic benefits of antibiotic treatment, contributing to the high recurrence rates observed in conditions like BV and the persistence of symptoms. However, alternatively, bEVs derived from antibiotics-exposed bacterial culture may enhance local inflammation in a manner that aids in the clearance of more pathogenic organisms. For instance, pre-incubation of LC with VK2 cells prior to *Candida albicans* infection (the causative agent of vulvovaginal candidiasis) enhanced the immune response by increasing levels of IL-2 (Th1-immunoregulatory), IL-6 (pro-inflammatory), and IL-17 (Th17 responses), while reducing IL-8 (chemotactic cytokine) ^29^. Understanding that LC bEVs may play a key role in mediating the beneficial effects of LC to the host, the alterations in their functionality, and the reduced bEV concentration following metronidazole exposure highlight the need for further and careful investigation. Understanding how antibiotic treatments intended to treat BV and other infections affect the functionality of bEVs from beneficial microbes is critical for optimizing therapeutic strategies that preserve or enhance host-microbiome homeostasis. Disruptions of this homeostatic state may be associated with poor recovery following metronidazole treatment.

Our analysis of human vaginal swabs confirmed the presence of bacterial GV bEVs in the vagina by targeting vaginolysin, which is uniquely produced by GV ^21^. Vaginolysin-EVs were detected even in subjects classified as CST I, despite a low relative abundance of GV. This aligns with previous findings showing that GV can be present in what is considered a healthy vaginal ecosystem, dominated by lactobacilli and characterized by a low Nugent score (0-3) ^30,31^. Our result suggests that taxonomic composition does not necessarily reflect the functional activity of the bacteria as previously reported by paired metagenomic and metatranscriptomic analyses ^32^. Even at low abundance, GV can remain transcriptionally active, producing bioactive molecules that can potentially be harmful.

The current study’s limitations include 1) bEVs were studied from single bacterial cultures, not from a polymicrobial environment; 2) no well-established normative data for produced bEVs in the human vagina; and 3) bEV isolation was conducted once per bacterial culture condition, without independent biological replicates. In addition, we did not include an unused swab control for ONI analysis, and we have not evaluated multiple strains of the same species, which have been show to co-exist in the vaginal microenvironment ^33^. BEVs generated from different bacterial strains may have distinctive roles in host-microbe interaction, as well as microbe-microbe crosstalk (*e*.*g*., inhibit the growth of certain species and support the growth of others while producing different metabolites affecting the host epithelium) ^34^. Furthermore, bEVs generated at different phases of the growth curve can have distinct cargo compositions ^35^, suggesting they possess different biological functions. Lastly, a recent clinical study found that oral and topical antimicrobial therapy in male partners of women with BV resulted in a significantly lower recurrence rate ^36^, highlighting the complexity of the treatment regimen and cautioning against generalized treatment. Investigating how antibiotics affect the cervicovaginal microbiome of affected women with BV and their partners, and the composition of bacterial products including bEVs by using proteomics approaches, could help further refine therapeutic approaches and reduce the risks of recurrent infections and adverse reproductive health outcomes.

In conclusion, our work reports that antibiotic treatments at sub-MIC alter the immunomodulatory properties of bEVs produced from the common vaginal anaerobes by altering the activity of bacterial recognition signaling pathways. This finding may provide a biological explanation for the high prevalence of symptoms after treatment for BV and/or for the high rate of recurrence of BV despite the adequacy of antibiotic treatment. Whether increased or decreased, an altered inflammatory profile from antibiotic-exposed bEVs may result in epithelial injury, unresolved inflammation, and/or promote continued vaginal dysbiosis. While antibiotic use is common and often indicated, the impact of these drugs on the microbiome and host-microbiome interactions requires further investigation. Taken together, these results suggest that alternative therapeutic approaches, such as probiotics and/or vaginal microbiota transplantation, may ultimately prove more effective and potentially reduce recurrence risk; however, future studies are needed to rigorously evaluate the efficacy and mechanisms of such strategies. The results herein argue for a better understanding of how antibiotic treatments for both vaginal and non-vaginal conditions may adversely impact the cervicovaginal microbiome and thus engender suboptimal reproductive health.

## Ethics approval and consent to participate

Not applicable.

## Consent for publication

Not applicable.

## Availability of data and materials

All data generated or analyzed during this study are included in this article and its supplementary information file.

## Acknowledgements

Mount Sinai Human Immune Monitoring Center for Luminex data acquisition support.

## Author contributions

Conception and design of the work: YH, ME; data acquisition: YH, OS, UR, MF, LN, IM; data analysis, interpretation, and drafted the manuscript: YH; revisions: YH, IM, HZ, JR, ME; supervision: JR, ME. All authors read and approved the final manuscript.

