## Supplementary Information for "Exposure to Antibiotics Modifies the Immune Profiles of Bacterial Extracellular Vesicles from Common Vaginal Anaerobes"

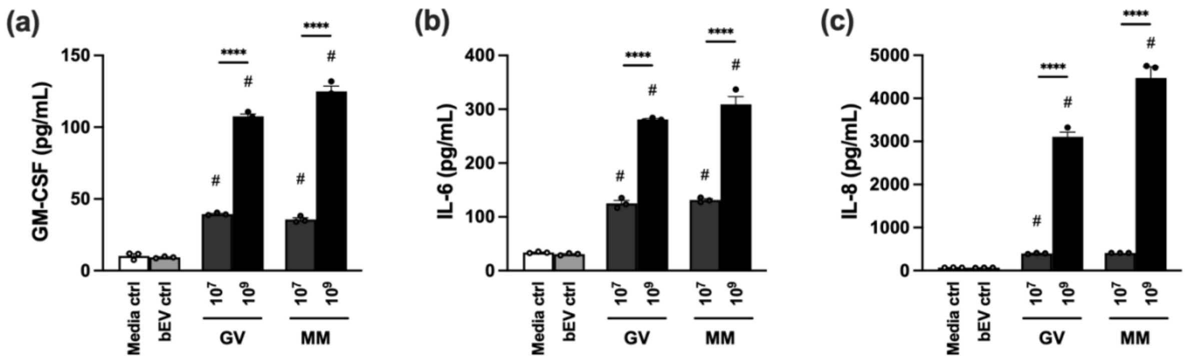


**Supplementary Figure 1.** The levels of inflammatory mediators identified by multiplex immunoassay in response to different doses of bEV treatment. (a) GM-CSF, (b) IL-6, and (c) IL-8 levels produced by End1 cells in response to $10^7$ or $10^9$ bEVs from Untreated GV or MM cultures (the results from $10^8$ are reported in the main text). Bar plots represent the mean, with error bars indicating the standard deviation (n=3). A one-way ANOVA followed by Sidak's multiple comparison test was used to compare group differences. Significant differences from the bEV vehicle control group are indicated with "#," and significant differences between bEV-treated groups are indicated with "" ($p < 0.05$), "" ($p < 0.01$), "" ($p < 0.001$), or "****" ($p < 0.0001$).

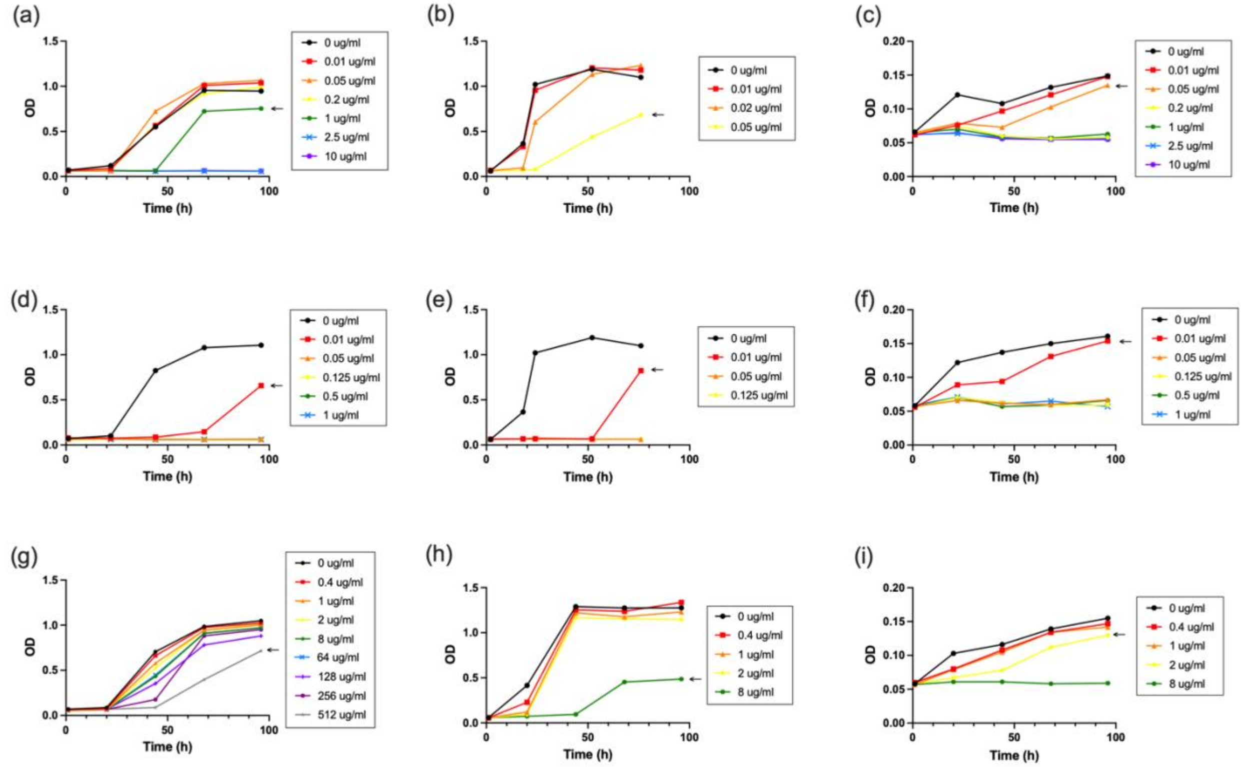


**Supplementary Figure 2.** OD measurements of LC, GV, and MM cultures with and without various doses of ABX over the course of 96 hours. (a) LC + Amp, (b) GV + Amp, (c) MM + Amp, (d) LC + Cli, (e) GV + Cli, (f) MM + Cli, (g) LC + Met, (h) GV + Met, (i) MM + Met. The sublethal MIC (Sub-MIC) used in the study is indicated by the black arrow in the figures.
